# CNTF specifically slows down the axonal transport of signalling endosomes

**DOI:** 10.1101/2025.10.09.681259

**Authors:** Elena R. Rhymes, James N. Sleigh, Giampietro Schiavo, Andrew P. Tosolini

## Abstract

Efficient axonal transport is essential for maintaining neuronal function, enabling the bidirectional delivery of diverse cargoes between the cell body and distal compartments. In the neuromuscular system, neurotrophic factors (NTFs) regulate motor neuron survival, function, and synaptic connectivity, in part, through retrograde trafficking of activated NTF-receptor complexes from the neuromuscular junction (NMJ) to the cell body. We recently demonstrated that brain-derived neurotrophic factor (BDNF) stimulation to muscles selectively enhances retrograde transport of signalling endosomes in fast, but not slow, motor neurons *in vivo*. Moreover, both axonal endosome transport and its BDNF-mediated regulation are disrupted in mouse models of diseases impacting motor neurons. Here, we examined whether additional NTFs, when applied to distal axon terminals, share this transport-modulating property. Through imaging sciatic nerves in anaesthetised mice, we tracked the *in vivo* dynamics of signalling endosomes in fast (FMN) and slow motor neurons (SMN) via intramuscular injections of a fluorescently conjugated, atoxic fragment of tetanus neurotoxin (H_C_T). H_C_T was co-administered with ciliary neurotrophic factor (CNTF), hepatocyte growth factor (HGF), neurturin (NRTN), or proBDNF - four growth factors with known effects on motor neurons. Compared to vehicle-treated controls, proBDNF, HGF, and NRTN, produced no detectable change in transport dynamics. In contrast, CNTF markedly reduced endosome speeds in both FMNs and SMNs, indicating remarkable selectivity of specific NTFs in the regulation of signalling endosome transport in motor neurons. Understanding this selectivity may aid the development of muscle-targeted NTF-based therapeutic strategies aimed at restoring axonal transport in neurodegenerative disease, peripheral neuropathy, and nerve injury.

**Graphical Abstract:** 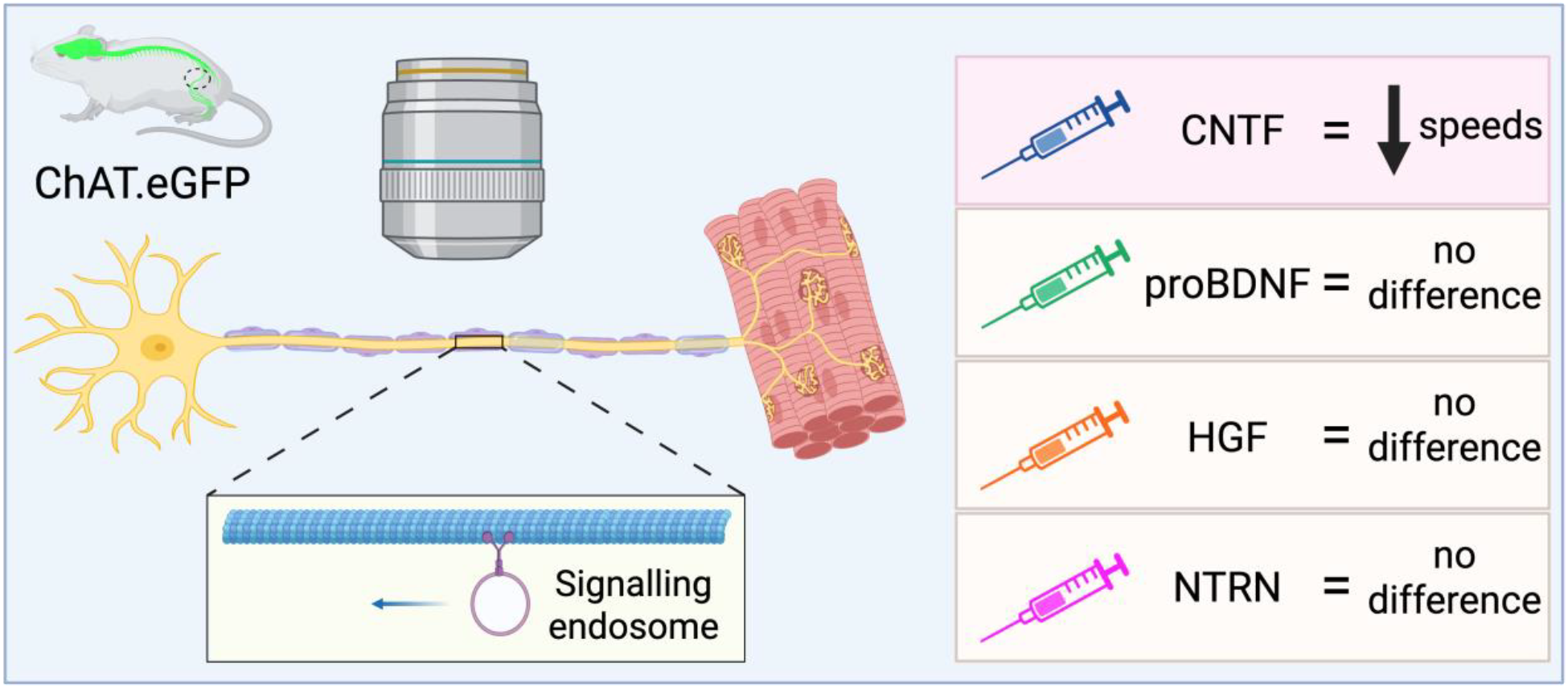

## Introduction

Neurotrophic factors (NTFs) are essential growth factors that regulate the development, maintenance, and plasticity of the central and peripheral nervous systems, including the neuromuscular synapse. During embryogenesis, NTFs, including ciliary neurotrophic factor (CNTF), brain-derived neurotrophic factor (BDNF), and neurotrophin-3 (NT-3) guide the survival, differentiation, and target innervation of cholinergic motor neurons (Wong et al., 1993). At the neuromuscular junction (NMJ), which is the specialised synapse formed between lower motor neuron termini and skeletal muscle fibres, bidirectional neurotrophin signalling events support formation, maturation and refinement of the synapse (Lu & Je, 2003; Chevrel et al., 2006; Pitts et al., 2006). However, rather than acting uniformly, NTFs exert distinct, context-dependent effects on the NMJ, reflecting a high degree of functional specificity. For example, BDNF and NT-3, but not nerve growth factor (NGF), enhance synaptic activity during development (Lohof et al., 1993). Additionally, differences in NTF processing and maturation (e.g., proBDNF vs mature BDNF) can determine receptor and co-receptor binding, further diversifying the influence of NTFs on synaptic and muscle physiology (Yang et al., 2009; Sakuma & Yamaguchi, 2011).

In adulthood, NTFs and their receptors adapt to the functional demands of the mature nervous system, assuming regulatory roles in synaptic transmission, neuronal excitability, plasticity and regenerative capacity (Wit et al., 2006; Gordon, 2009; Dorsey et al., 2011; Pérez et al., 2019; Yanpallewar et al., 2021). Teasing apart the embryonic versus mature, and context-specific roles of neurotrophic signalling is essential for understanding how these factors shape motor neuron physiology throughout development and into adulthood (Tovar-y-Romo et al., 2014), and their potential therapeutic relevance in neuromuscular disorders.

Motor units, comprising α-motor neurons and the skeletal muscle fibres they innervate, are broadly categorised as fast or slow. Fast motor units couple fast motor neurons (FMNs) with fast-twitch type IIb (glycolytic, fatigable), IIx, and IIa (oxidative-glycolytic, fatigue-resistant) muscle fibres to produce strong, rapid muscle contractions, whereas slow motor units pair slow motor neurons (SMNs) with slow-twitch type I (oxidative, fatigue-resistant) fibres for sustained, low-force activity (Stifani, 2014; Suetterlin et al., 2022). These functional differences are mirrored by distinct motor neuron properties, including size, excitability, metabolic profiles, and gene expression (Burke et al., 1973; Kanning et al., 2010; Blum et al., 2021).

Emerging evidence indicates that NTFs not only support motor neuron survival but also motor unit identity (Stansberry & Pierchala, 2023). Indeed, BDNF within skeletal muscle promotes glycolytic fibre specification (Delezie et al., 2019), whereas muscle-secreted neurturin (NRTN) enhances SMN identity (Correia et al., 2021), and such subtype-specific regulation may underl differential responsiveness to NTF signalling. Notably, increasing the availability of BDNF at motor nerve terminals enhances axonal transport dynamics selectively in FMNs, which exhibit reduced sensitivity to BDNF and impaired cargo trafficking in neurodegenerative conditions and with advanced age (Shekari & Fahnestock, 2019; Tosolini et al., 2022; Sleigh et al., 2023; Rhymes et al., 2024; Villarroel-Campos et al., 2025a).

Given the reliance of long-range NTF signalling on intracellular trafficking, axonal transport is central to motor neuron communication and survival (Wu et al., 2009). This microtubule-based process enables neurons to shuttle cargo over long distances, with retrograde transport being crucial for conveying trophic signals from the NMJ to the soma. The signalling endosome serves as the principal vehicle for retrograde transport of NTF-receptor complexes from the NMJ to the motor neuron soma (Villarroel-Campos et al., 2018), and its dynamics can be assessed in both *in vivo* and *in vitro* experimental systems with the application of live fluorescent imaging (Surana et al., 2020). The compartmentalised nature of NTF-receptor interactions suggests that neurotrophins exert spatially distinct effects across motor neuron subtypes. Pioneering this concept, DiStefano et al., (1992) demonstrated that BDNF, NT-3, and NGF undergo distinct patterns of retrograde transport in central and peripheral neurons. More recently, Schaller et al., (2017) showed that individual NTFs selectively support the survival of discrete embryonic motor neuron subtypes (e.g., CNTF binding to LIFRβ/gp130 for axial motor neurons; hepatocyte growth factor (HGF) binding to mesenchymal-epithelial transition factor (c-Met) for limb motor neurons; and artemin binding to GFRα3-RET for pelvic motor neurons), an effect that exhibited additive impact when applied in combination. These findings emphasise the specificity of NTF signalling and the importance of dissecting their roles in intracellular trafficking and long-range signal propagation in motor neurons.

To assess whether distal neurotrophic cues differentially influence axonal transport across motor neuron subtypes, we selected CNTF, proBDNF, HGF, and NRTN for their well-established roles in motor neuron physiology and regeneration. CNTF was selected not only for its function in promoting axonal regeneration (Bauer et al., 2007), but also due to its widespread use in neuronal cultures to enhance motor axon growth and survival (Schaller et al., 2017; Rhymes et al., 2022; Tosolini et al., 2022). Expanding upon our previous reports that FMN transport is enhanced following BDNF administration (Tosolini et al., 2022), we selected proBDNF to investigate the effect of specifically engaging p75^NTR^ and sortilin, which are considered to drive largely negative/ regressive signalling events (Lalli & Schiavo, 2002; Butowt & Bartheld, 2009). HGF is a pleiotropic growth factor that can be secreted from muscle and promotes embryonic motor neuron survival, with synergistic effects when combined with other NTFs (Yamamoto et al., 1997; Schaller et al., 2017). Finally, NRTN was chosen for its established role in promoting SMN identity, and to tease apart motor neuron subtype-specific effects of diverse NTFs (Correia et al., 2021). Accordingly, the receptors for these NTFs have been observed in skeletal muscle, including at the NMJ. These include CNTFRα, LIFR, and gp130 for CNTF (Guillet et al., 1999), RET for NRTN signalling complexes (Baudet et al., 2008), the HGF receptor c-Met (Lindström et al., 2010; Webster & Fan, 2013), and the proBDNF receptors p75^NTR^ and sortilin (Ariga et al., 2017; Pérez et al., 2019; Tosolini et al., 2022).

Using a powerful intravital imaging platform to assess the axonal transport of signalling endosomes in an intact and mature neuromuscular system (Sleigh et al., 2017), we evaluated whether these four NTFs acutely modulate transport dynamics when delivered to motor nerve terminals. We found that only CNTF influences axonal transport dynamics and did so by reducing mean endosomal speeds. These insights deepen our understanding of motor unit biology and provide a foundation for modulating distal neurotrophin signalling in disease and/or ageing contexts.

## Materials and Methods

### Ethical Approval

All animal procedures were conducted in accordance with the UK Animals (Scientific Procedures) Act (1986) under a license issued by the UK Home Office and were approved by the UCL Queen Square Institute of Neurology Ethical Review Committee.

### Animals

Tg(Chat-EGFP)GH293Gsat/Mmucd mice (RRID: MMRRC_000296-UCD), hereafter referred to as ‘ChAT.eGFP’ mice, were maintained in heterozygosity on a CD-1 background, as previously described (Sleigh et al., 2020a; Tosolini et al., 2024). Animals were housed in individually ventilated cages under controlled temperature and humidity conditions, with a 12 h light/dark cycle and free access to food and water. Experimental cohorts included both males (n = 17) and females (n = 22), aged ∼2-3.5 months (mean = 90 days), an interval during which axonal transport dynamics in wild-type mice are known to remain consistent (Sleigh et al., 2020a, 2023). All procedures and animal care protocols were conducted in accordance with the ARRIVE (Animal Research: Reporting of *In Vivo* Experiments) guidelines to ensure ethical and reproducible research standards.

### In vivo axonal transport imaging

Retrograde axonal transport of signalling endosomes was visualised using an atoxic fragment of tetanus neurotoxin (H_C_T; residues 875–1315), fused to a cysteine-rich domain and a human influenza HA epitope and labelled with AlexaFluor555 C2 maleimide (Thermo Fisher Scientific, A-20346), using reported protocols (Lalli & Schiavo, 2002). To adhere to the 3Rs principles and to allow direct comparisons between FMNs and SMNs, H_C_T was injected under isoflurane-induced anaesthesia into the left tibialis anterior and the right soleus muscles of the same animal (**Fig. 1B**), as previously described (Gibbs et al., 2016; Sleigh et al., 2020b; Tosolini et al., 2021). Mice were divided into five experimental cohorts, and received 5-7 µg of H_C_T mixed with: a) phosphate buffered saline (PBS) as the control; b) 50 ng recombinant human CNTF protein (Peprotech, 450-13), c) 50 ng recombinant mouse cleavage-resistant proBDNF (Alomone labs, B-243), d) 25 ng recombinant human HGF (Bio-Techne, 294-HG[CF]), or e) 50 ng recombinant human NTRN (Peprotech, 450-11), as summarised in **Figure 1A**. 4-8 h later, mice were re-anaesthetised, and the first sciatic nerve was exposed for imaging. Time-lapse imaging was performed on a Zeiss LSM 780 confocal microscopes equipped with 40× Plan-Apochromat oil immersion objective encased within an environmental chamber maintained at 37°C. Images were acquired at ∼1 s intervals. At the completion of the imaging of the first sciatic nerve, the contralateral sciatic nerve was exposed and imaged. To minimise bias, the order of imaging between the left (tibialis anterior-injected) and right (soleus-injected) sciatic nerves was randomised. eGFP-positive cholinergic motor axons were imaged, and regions containing nodes of Ranvier were avoided (Sleigh et al., 2020a; Tosolini et al., 2024). Following imaging of both sciatic nerves, animals were euthanised within 1 h of terminal anaesthesia induction.

**Figure 1.**
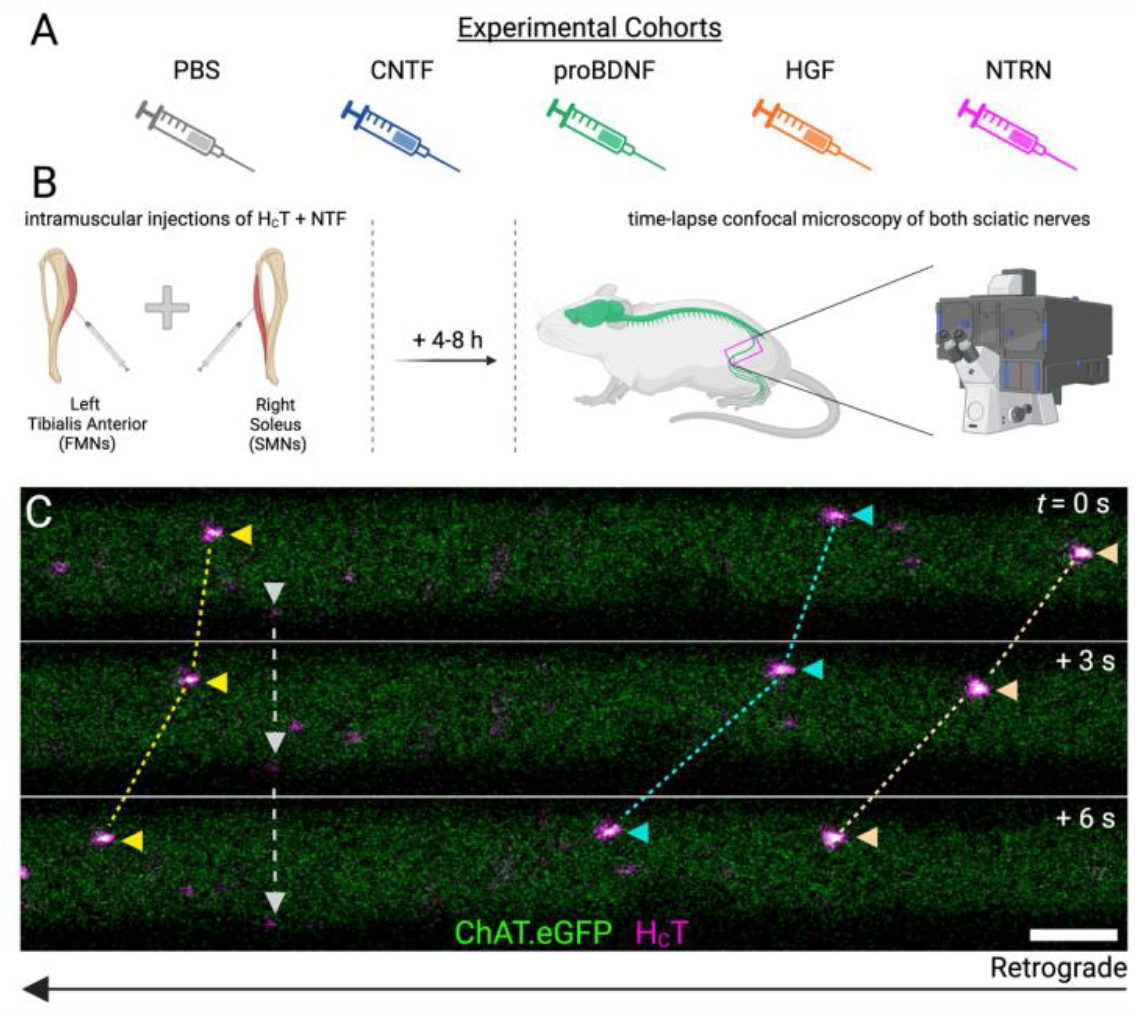
Experimental design. **A)** Male and female ChAT.eGFP mice were allocated into one of five groups: *i)* PBS (control); *ii)* CNTF; *iii)* proBDNF; *iv)* HGF; and *v)* NTRN. **B)** Each mouse received intramuscular injections of H_C_T combined with a single NTF into the left tibialis anterior and right soleus muscles to target fast and slow motor neurons, respectively. After a 4-8 h incubation period, time-lapse microscopy was performed on both sciatic nerves. **C)** H_C_T-labelled signalling endosomes (pseudo-coloured in magenta) from single ChAT.eGFP motor axons were individually tracked to quantify retrograde transport dynamics. Three representative retrogradely transported signalling endosomes are identified by yellow, cyan, and peach arrowheads connected by dashed lines across frames. Grey arrowheads and dashed lines identify a stationary endosome. See also **Video 1**. Scale bar = 5 μm, frame interval = 3 s.

### In vivo axonal transport analysis

A minimum of 15 endosomes per axon from at least three axons per mouse were manually tracked using the FIJI plugin ‘TrackMate‘ (**Fig. 1C**; **Video 1**) (Ershov et al., 2022). Only retrogradely moving signalling endosomes that could be tracked for ≥ 10 consecutive frames were included. An endosome was classified as paused if it remained within a ≈ 0.2 μm interval for at least two consecutive frames (2 s) to account for minor displacement caused by breathing artefacts. Endosomes were excluded from the analysis if they were immobile for ≥ 10 consecutive frames (≈ 20 s), or failed to meet the minimum track duration of (*i.e*., they moved for ≤ 10 consecutive frames). The “pausing %” was calculated per animal by determining the number of frames in which endosomes were paused, dividing this by the total number of frames analysed and multiplying this value by 100.

### Statistical Analysis

All statistical analyses were performed at the conclusion of the tracking. Data were assessed with two-way analyses of variance (ANOVA) with subsequent Holm-Šídák^1^s multiple comparisons tests. Sample sizes were pre-determined using power calculations and previous experience (Rhymes et al., 2022, 2024; Tosolini et al., 2022; Sleigh et al., 2023; Simkin et al., 2025). Means ± standard error of the mean (SEM) are plotted for all graphs. GraphPad Prism 10 software (version 10.5.0) was used for statistical analyses and figure production.

## Results

### In Vivo Axonal Transport of Signalling Endosomes is Similar in FMNs and SMNs and Between Sexes

Our intravital imaging platform enables real-time visualisation of retrograde axonal transport of neurotrophin-containing signalling endosomes using muscle-injected fluorescent H_C_T (Gibbs et al., 2016; Sleigh et al., 2020b; Tosolini et al., 2021) (**Video 1**). By injecting selected muscles from contralateral hindlimbs in ChAT.eGFP mice, we compared signalling endosome transport dynamics between FMNs and SMNs within the same animal (**Figure 1**). To minimise age-related variability (Milde et al., 2015; Villarroel-Campos et al., 2025a), all experiments were performed in mice aged approximately 3 months (mean = 90 days).

We first wanted to establish whether baseline transport parameters were similar between sexes and in motor neuron subtypes. H_C_T was therefore injected into the left tibialis anterior (TA) and right soleus (Sol) muscles, two muscles previously shown to exhibit comparable transport dynamics under basal conditions, yet divergent responses to BDNF stimulation and transport impairments in ALS models (Tosolini et al., 2022, 2024). We observed that males and females exhibit similar signalling endosome dynamics in FMNs and SMNs, with no differences in mean speed, maximum speed or pausing events (**Figure 2**). Therefore, to adhere to the 3R’s principles and minimise animals used in the following experiments, data from males and females were combined.

**Figure 2.**
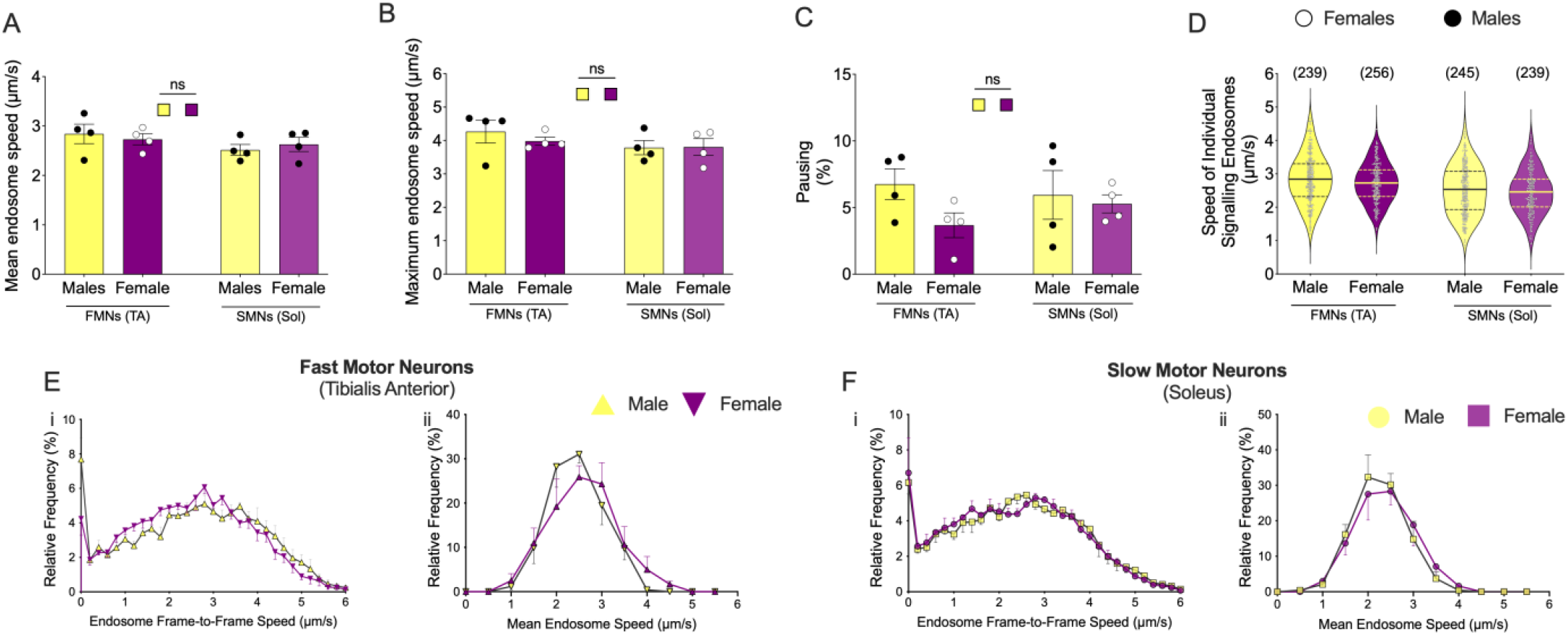
Under basal conditions, the axonal transport dynamics of signalling endosomes are equivalent in male and female mice. In both fast motor neurons (FMNs) and slow motor neurons (SMNs), males and females exhibit comparable **A**) mean endosome speed (*p* = 0.17 for motor neuron type; *p* = 0.99 for sex; *p* = 0.46 for interaction), **B**) maximum endosome speed (*p* = 0.21 for motor neuron type; *p* = 0.60 for sex; *p* = 0.54 for interaction), and **C**) pausing percentage (*p* = 0.75 for motor neuron type; *p* = 0.15 for sex; *p* = 0.34 for interaction). **D**) Violin plots of individual endosomes also indicate similar speed profiles in FMNs and SMNs of both males and females (FMNs: male mean = 2.84 µm/s ± 0.05 [n = 239], female mean = 2.73 µm/s ± 0.04 [n = 256]; SMNs: male mean = 2.52 µm/s ± 0.05 [n = 245], female mean = 2.44 µm/s ± 0.04 [n = 239]). Furthermore, the endosome frame-to-frame and mean endosome speed distribution curves closely overlap between the sexes for **E**) FMNs and **F**) SMNs, indicating no sex-specific differences in baseline retrograde transport dynamics. Statistical analyses were performed using two-way ANOVA and Holm-Šídák^1^s multiple comparisons tests. ns, not significant. n = 4.

### CNTF Slows Retrograde Transport in Fast and Slow Motor Neurons

CNTF is a cytokine-like NTF that promotes motor neuron survival and axonal regeneration via the LIFRβ/gp130 receptor complex (Bauer et al., 2007). It is commonly applied *in vitro* to support axonal growth, can protect vulnerable motor neurons from early degeneration in ALS mice (Pun et al., 2006), can compensate against retrograde deficits observed in the *pmn* mouse model of ALS (Sagot et al., 1998) and can regulate muscle strength during ageing (Guillet et al., 1999). However, the direct effect of CNTF on axonal transport has not been fully elucidated. To investigate this, we co-injected 50 ng of recombinant CNTF and H_C_T into the left TA and right Sol, followed by intravital imaging of both sciatic nerves 4-8 h later (**Figure 1**). Given, its overt pro-survival role, we hypothesised that CNTF supplementation would increase transport speeds. Unexpectedly, we found that CNTF did the opposite by reducing mean endosome speeds in both FMNs and SMNs, without influencing maximum speed or pausing (**Figure 3A-D**). This was further supported by a leftward shift in frame-to-frame and mean speed distributions, indicating a global slowing of retrograde transport (**Figure 3E-F**). These observations provide the first evidence that CNTF can negatively modulate, rather than enhance, endosomal transport dynamics, adding a new dimension to its established roles in motor neuron biology.

**Figure 3.**
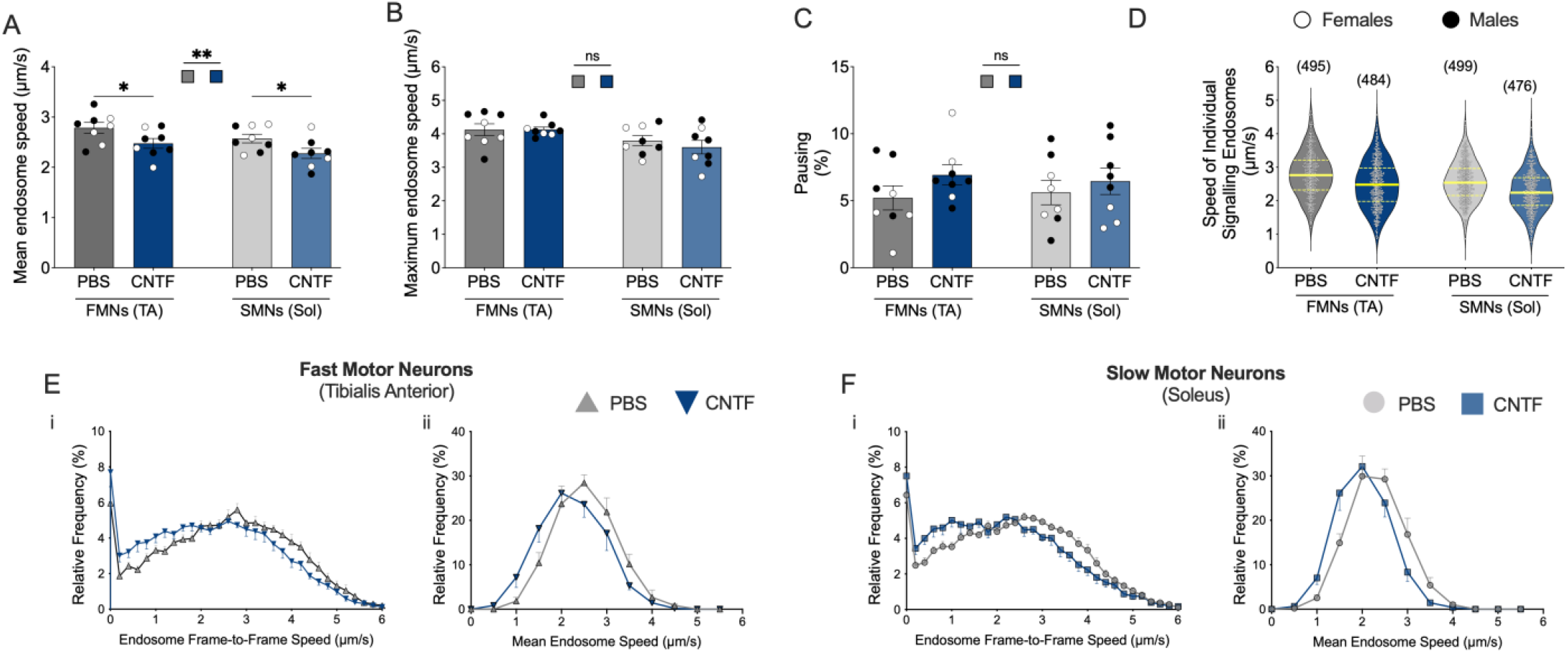
Acute treatment of muscles with CNTF reduces mean signalling endosome transport speed in both fast motor neurons (FMNs) and slow motor neurons (SMNs). **A**) Acute intramuscular injection of CNTF significantly reduces the mean endosome speed in both FMNs and SMNs (*p* = 0.10 for motor neuron type; *p* = 0.001 for stimulation factor; *p* = 0.92 for interaction). Further, this effect is not associated with differences in the **B**) maximum endosome speed (*p* = 0.05 for motor neuron type; *p* = 0.40 for stimulation factor; *p* = 0.38 for interaction) or **C**) pausing percentage (*p* = 0.97 for motor neuron type; *p* = 0.18 for stimulation factor; *p* = 0.64 for interaction). **D**) Violin plots of individual endosome speeds show slower transport in CNTF-treated mice (FMNs: PBS mean = 2.78 µm/s ± 0.03 [n = 495], CNTF mean = 2.48 µm/s ± 0.03 [n = 484]; SMNs: PBS mean = 2.57 µm/s ± 0.03 [n = 495], CNTF mean = 2.27 µm/s ± 0.03 [n = 476]). The leftward shift in both frame-to-frame and mean endosome speed distributions for CNTF-treated **E**) FMNs and **F**) SMNs, indicate slower retrograde transport following CNTF exposure. Statistical analyses were performed using two-way ANOVA and Holm-Šídák^1^s multiple comparisons tests. * < 0.05, ** < 0.01. ns, not significant. n = 8. Black circles = males (n = 4 PBS; n = 5 CNTF), white circles = females (n = 4 PBS; n = 3 CNTF).

### proBDNF Does Not Influence Retrograde Endosomal Movement in FMNs or SMNs

proBDNF is the precursor to mature BDNF and signals primarily through the p75^NTR^-sortilin receptor complex, driving processes such as synaptic pruning and apoptosis (Lu et al., 2005). While mature BDNF elicits pro-survival signalling and selectively enhances axonal transport in motor neurons (Tosolini et al., 2022), the effects of proBDNF remains unclear. To address this, we injected 50 ng of recombinant, cleavage-resistant proBDNF along with H_C_T into the left TA and right Sol muscles, followed by intravital imaging of the corresponding sciatic nerves 4-8 h later (**Figure 1**). We speculated that proBDNF might negatively influence axonal transport through engaging p75^NTR^-sortilin complexes. In contrast, proBDNF stimulation did not alter mean endosome transport speeds, maximum velocities, or pausing events across both FMNs and SMNs (**Figure 4A-D**). Distribution curves of frame-to-frame and mean endosome speeds were largely overlapping with control (**Figure 4E-F**). Collectively, these data indicate that proBDNF does not modulate retrograde endosomal transport in healthy mice in either motor neuron subtypes.

**Figure 4.**
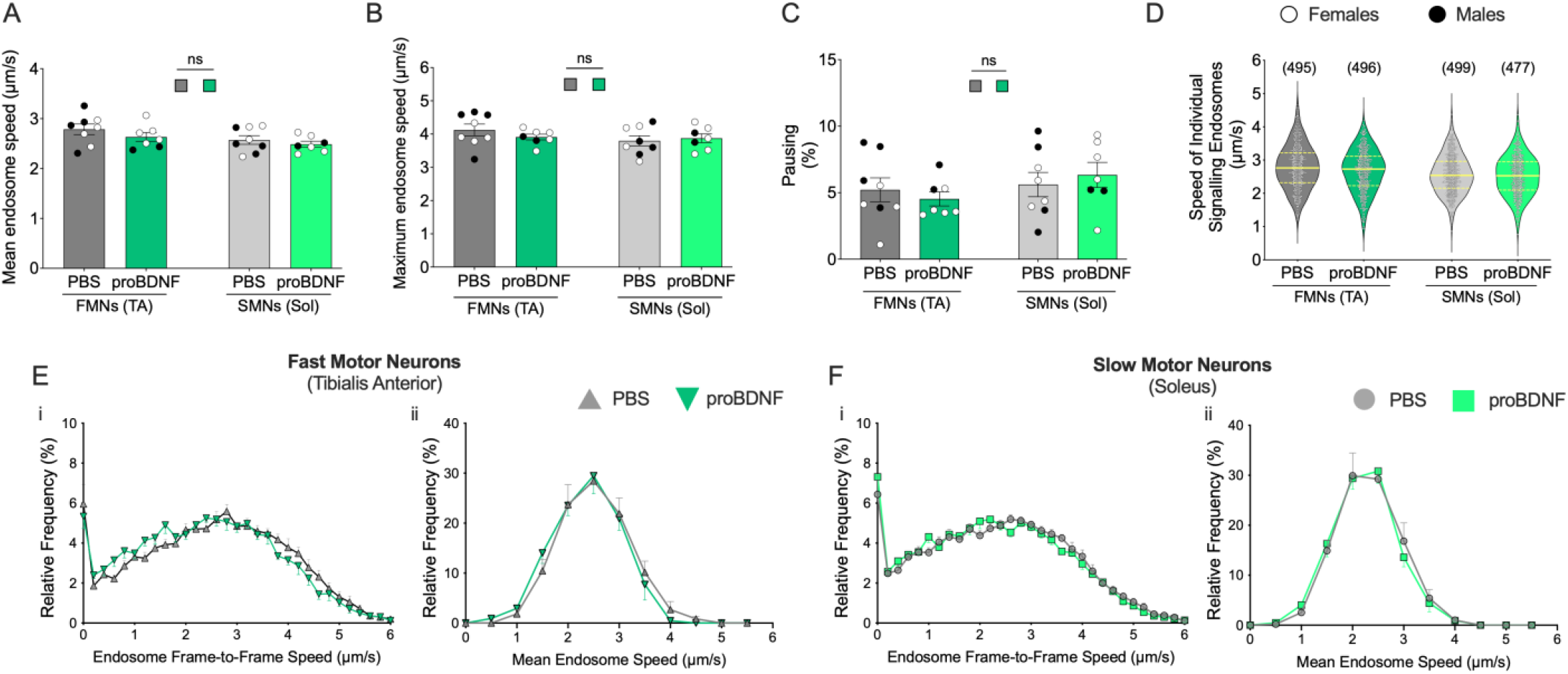
Stimulation of muscles with proBDNF does not influence the axonal transport of signalling endosome. In both fast motor neurons (FMNs) and slow motor neurons (SMNs), proBDNF did not alter **A)** mean endosome speed (*p* = 0.06 for motor neuron type; *p* = 0.19 for stimulation factor; *p* = 0.70 for interaction), **B)** maximum endosome speed (*p* = 0.30 for motor neuron type; p = 0.62 for stimulation factor; p = 0.25 for interaction) or **C)** pausing percentage (*p* = 0.24 for motor neuron type; *p* = 0.92 for stimulation factor; *p* = 0.34 for interaction). **D**) Violin plots of individual endosomes show comparable distributions in all conditions (FMNs: PBS mean = 2.78 µm/s ± 0.03 [n = 495], proBDNF mean = 2.67 µm/s ± 0.03 [n = 496]; SMNs: PBS mean = 2.57 µm/ s ± 0.03 [n = 495], proBDNF mean = 2.53 µm/s ± 0.03 [n = 477]). Overlapping endosome frame-to-frame and mean endosome speed distribution curves confirm that proBDNF does not modulate retrograde transport in **E)** FMNs or **F)** SMNs. Statistical analyses were performed using two-way ANOVA and Holm-Šídák^1^s multiple comparisons tests. ns, not significant. n = 7-8. Black circles = males (n = 4 PBS; n = 2 CNTF), white circles = females (n = 4 PBS; n = 5 proBDNF).

### Axonal Transport is not modulated by HGF

HGF, which acts through the c-MET receptor, serves as both a muscle-derived survival factor and chemoattractant during development, with pronounced effects on cervical and lumbar motor neurons innervating limb musculature (Yamamoto et al., 1997). Beyond its individual effects, HGF has been shown to ‘amplify’ the survival and regenerative responses of embryonic motor neurons when combined with other NTFs, including BDNF, CNTF and NRTN (Schaller et al., 2017). In the adult CNS, muscle-derived HGF may act as a retrograde survival factor, supporting cholinergic motor neurons after axonal injury (Okura et al., 1999). To test whether HGF modulates retrograde axonal transport, we co-injected 25 ng of recombinant HGF with H_C_T into the left TA and right soleus muscles, followed by intravital imaging on both sciatic nerves 4-8 h later (**Figure 1**). Contrary to our expectations, HGF had no effect on endosomal transport in either fast or motor neurons, with no change in mean speed, peak velocity, or pause frequency (**Figure 5A-D**). In addition, no notable changes were observed in the distribution curves of frame-to-frame or mean endosome speeds compared to controls (**Figure 5E-F**). These results indicate that acute muscle exposure to HGF does not regulate long-range axonal transport kinetics of signalling endosomes in both motor neuron subtypes.

**Figure 5.**
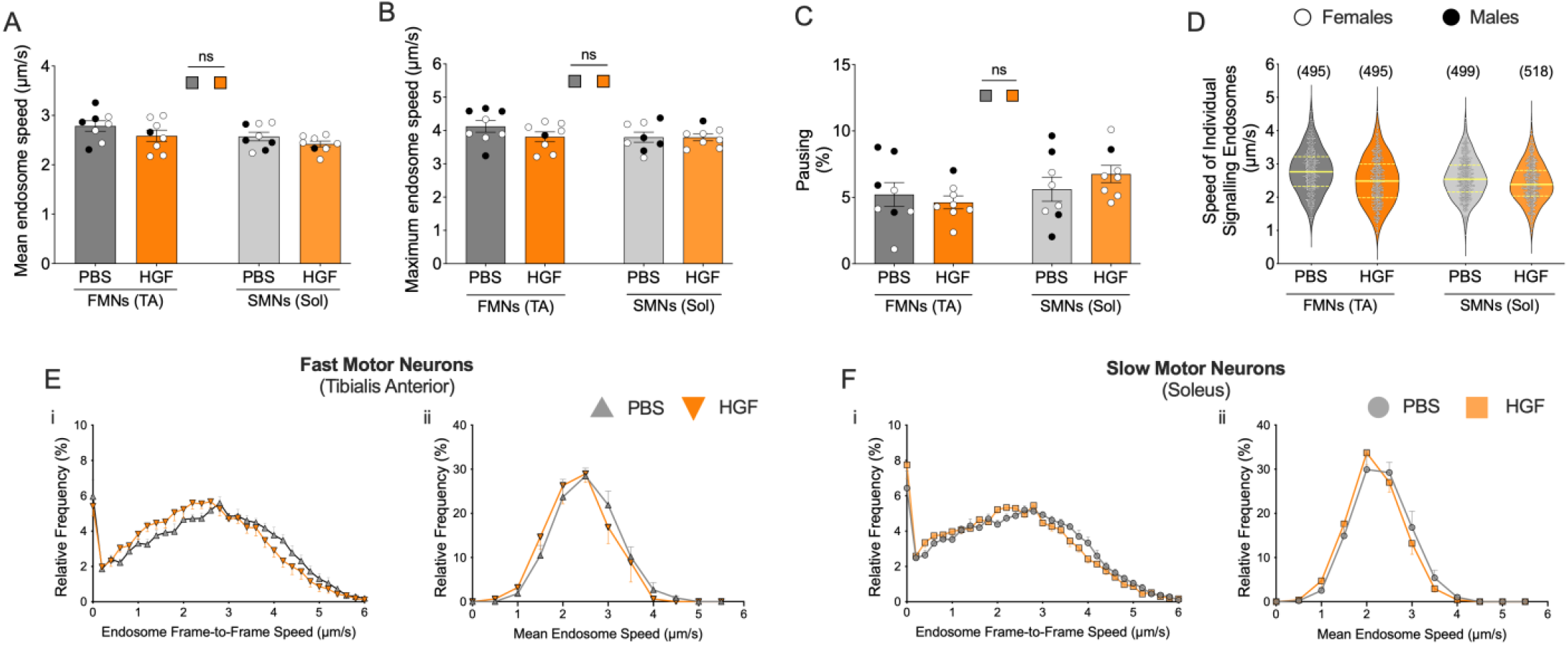
Stimulation of muscles with HGF does not affect retrograde transport of signalling endosomes. Following acute HGF treatment of muscles, no significant changes were observed in endosome transport parameters across fast (FMNs) and slow (SMNs) motor neurons. Specifically, there were no differences in **A)** mean endosome speed (*p* = 0.02 for motor neuron type; *p* = 0.13 for stimulating factor; *p* = 0.83 for interaction), **B)** maximum endosome speed (*p* = 0.14 for motor neuron type; *p* = 0.38 for stimulating factor; *p* = 0.40 for interaction), or **C)** pausing frequency (*p* = 0.12 for motor neuron type; *p* = 0.72 for stimulating factor; *p* = 0.27 for interaction). **D)** Violin plots of individual endosome speeds further demonstrate the similarity between PBS- and HGF-treated motor neurons (FMNs PBS mean = 2.78 µm/s ± 0.03 [n = 495], HGF mean = 2.49 µm/s ± 0.03 [n = 496]; SMNs mean = 2.57 µm/s ± 0.03 [n = 495], HGF mean = 2.42 µm/s ± 0.03 [n = 477]). Distribution plots for both frame-to-frame and mean endosome speeds overlap substantially between HGF and control groups, indicating no modulation of retrograde transport by HGF in **E)** FMNs and **F)** SMNs. Statistical analyses were performed using two-way ANOVA and Holm-Šídák^1^s multiple comparisons tests. ns, not significant. n = 8. Black circles = males (n = 4 PBS; n = 1 HGF), white circles = females (n = 4 PBS; n = 7 HGF).

### Axonal Transport is Unaffected by Short-Term NTRN Exposure

NRTN is a member of the GDNF family of ligands that signals through a receptor complex composed of GFRα2 and the tyrosine kinase RET to support the survival and maintenance of neurons (Golden *et al*., 1998). At the NMJ, NRTN influences both pre- and post-synaptic compartments by promoting motor axon growth, enhancing synaptic vesicle accumulation, and stimulating acetylcholine receptor clustering on muscle fibres (Wang *et al*., 2002; Yang & Nelson, 2004). More recently, NRTN was identified to be a PGC-1α1-regulated myokine, secreted from skeletal muscle that retrogradely signals to motor neurons to support NMJ formation, motor neuron recruitment, and the transcriptional program regulating SMN identity (Mills *et al*., 2018; Correia *et al*., 2021). To determine whether NRTN directly modulates axonal transport, we co-injected 50 ng of recombinant NRTN with H_C_T into the left TA and right soleus muscles and performed intravital imaging of sciatic nerves 4-8 h later. Despite its role in shaping SMN identity, NRTN had no detectable effects on retrograde transport dynamics; with endosome speeds, velocity distributions, and pause frequencies remaining comparable to controls (**Figure 6**). These results suggest that NRTN does not acutely modulate axonal trafficking *in vivo* in mature motor neurons.

**Figure 6.**
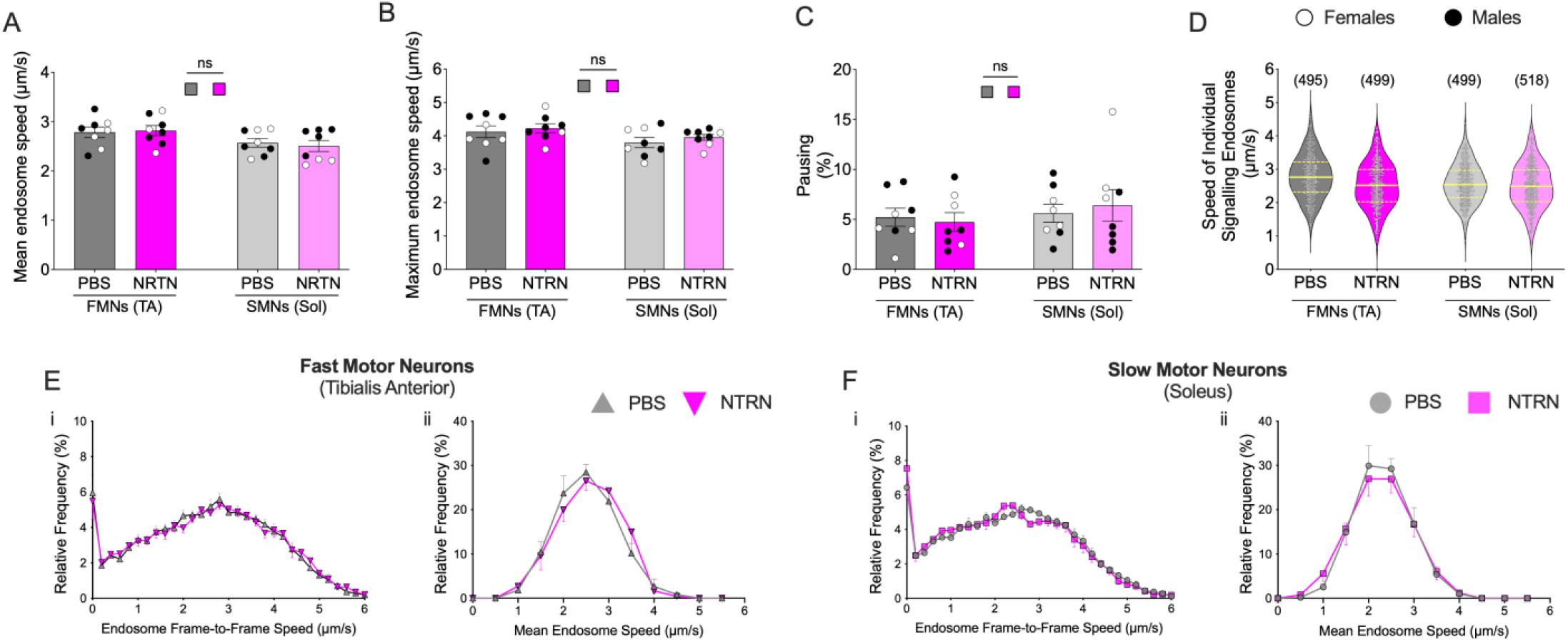
Stimulation of muscles with NRTN does not alter signalling endosome transport dynamics. In both fast (FMNs) and slow (SMNs) motor neurons, acute stimulation of muscles with NRTN resulted in no detectable changes in endosomal trafficking dynamics. In both FMNs and SMNs, NTRN stimulation did not influence **A)** mean endosome speeds (*p* = 0.02 for motor neuron type; *p* = 0.88 for stimulating factor; *p* = 0.63 for interaction), **B)** maximum endosome speeds (*p* = 0.06 for motor neuron type; *p* = 0.36 for stimulating factor; *p* = 0.85 for interaction) or **C)** pausing percentages (*p* = 0.47 for motor neuron type; *p* = 0.85 for stimulating factor; *p* = 0.41 for interaction). D) Violin plots of individual endosome velocities confirmed that NRTN treatment did not alter transport profiles (FMNs PBS mean = 2.78 ± 0.03 [n = 495], NRTN mean = 2.51 ± 0.03 [n = 499]; SMNs mean = 2.57 ± 0.03 [n = 495], NRTN mea n = 2.50 ± 0.03 [n = 518]). Frame-to-frame and mean endosome speed distribution curves also largely overlapped between PBS- and NRTN-treated **E)** FMNs and **F)** SMNs, indicating no impact of NRTN on retrograde transport. Statistical analyses were performed using two-way ANOVA and Holm-Šídák^1^s multiple comparisons tests. ns, not significant. n = 8. Black circles = males (n = 4 PBS; n = 5 NRTN), white circles = females (n = 4 PBS; n = 3 NRTN).

## Discussion

NTFs are well-established regulators of motor neuron survival, identity, and neuromuscular connectivity. Our previous work demonstrated that BDNF selectively enhances signalling endosome transport in FMNs, but not SMNs, in healthy adult mice (Tosolini et al., 2022). Here, we expanded our analysis to additional NTFs with unknown *in vivo* effects on the axonal transport of signalling endosomes in motor neurons. Surprisingly, none of the four tested NTFs enhanced retrograde transport speeds; proBDNF, HGF, and NRTN showed no overt effects, whereas CNTF significantly slowed down endosome transport in both FMNs and SMNs. These findings reveal that the modulatory capacity on axonal transport is not a universal feature of NTFs and may depend, at least in part, on both the ligand and motor neuron subtype.

### Mechanistic Insights from BDNF

Much of what is known about NTF regulation of intracellular trafficking is derived from studies on BDNF, which has been shown to upregulate endocytosis, enhance endosomal flux, increase retrograde transport speeds, and subsequently drive transcriptional and translational changes in the soma (Roux et al., 2006; Wang et al., 2016; Tosolini et al., 2022; Moya-Alvarado et al., 2023; Sleigh et al., 2023; Rhymes et al., 2024). Importantly, the pro-survival actions of NTFs depend not simply on their presence in the soma, but on their successful delivery as activated receptor-ligand complexes from the axon (Heerssen et al., 2004).

This process begins at the distal axon, where BDNF-TrkB engagement activates ERK1/2 and AKT pathways and induces Ca^2+^ influx (Dombert et al., 2017; Moya-Alvarado et al., 2024), while Rab10 contributes to the sorting of TrkB into signalling endosomes that propagate retrograde signalling (Lazo & Schiavo, 2023). We recently identified that BDNF-dependent activation of ERK1/2 enhances axonal transport by modulating microtubule-associated protein phosphorylation, whereas AKT primarily supports survival signals within the soma (Vargas et al., 2025). Indeed, inhibition of axonal AKT does not impair transport and can even enhance it, underscoring that not all growth factor activated pathways promote faster transport (Fellows et al., 2020). Consistent with this, BDNF-driven enhancement of trafficking is both transient, dissipating within 24 hours, and ERK1/2-dependent, as sequestration of muscle-derived BDNF or inhibition of ERK1/2 impairs transport (Sleigh et al., 2023).

At the transport machinery level, Trk receptors directly associate with cytoplasmic dynein components, including the 14 kDa light chain (*i.e*., DLC) and the 74 kDa intermediate chain (*i.e*., DIC) (Yano et al., 2001). ERK1/2-mediated phosphorylation of DIC facilitates dynein recruitment to signalling endosomes (Mitchell et al., 2012), while neurotrophin signalling promotes synthesis of cofactors such as Lis1 and p150Glued to fine-tune cargo handling (Villarin et al., 2016). In parallel, BDNF-induced Ca^2+^ influx activates calcineurin, dephosphorylating huntingtin and initiating retrograde trafficking (Scaramuzzino et al., 2022). In BDNF-containing endosomes, glyceraldehyde-3-phosphate dehydrogenase (GAPDH) tethered to the vesicle surface generates ATP locally, supplying the energy required to sustain long-distance transport (Zala et al., 2013). Collectively, this represents a tightly coordinated system integrating receptor signalling, motor activation, and local metabolic support.

### Divergent NTF Effects

Despite converging on common signalling pathways, NTFs vary greatly in their effects on axonal transport. CNTF now joins IGF1 (Fellows et al., 2020) and VEGF-165 (Sleigh et al., 2023) as negative regulators of signalling endosome motility. Conversely, proBDNF, HGF, and NRTN, similar to NT-3 and NT-4 (Sleigh et al., 2023), exert no detectable influence on motor neuron signalling endosome transport *in vivo*. The case for GDNF is more complex, showing no effect *in vivo* (Tosolini et al., 2022), but *in vitro* assays revealed enhanced motility following GDNF treatment, either alone or with GFRα1, although GFRα1 alone did not elicit an effect (Rhymes et al., 2022). Whether these NTFs produce additive effects on transport, as observed in embryonic motor neuron survival (Schaller et al., 2017), remains unresolved.

An important consideration is that regulation of endosomal axonal transport by NTFs may be dose-dependent, where insufficient local ligand concentration fails to induce signalling, and excessive stimulation can trigger aberrant receptor internalisation and feedback inhibition. Indeed, phosphoproteomic and transcriptomic analyses have shown that prolonged BDNF exposure upregulates several negative regulators of downstream signalling, supporting the concept of dose-dependent feedback control (Vargas et al., 2025). Furthermore, administration of 25 ng BDNF has previously been shown to modulate transport *in vivo* (Tosolini et al., 2022; Sleigh et al., 2023; Rhymes et al., 2024; Villarroel-Campos et al., 2025a), suggesting the existence of an optimal concentration window for effective NTF regulation of trafficking, although this requires experimental validation.

Although both BDNF and CNTF activate ERK1/2, their opposing transport phenotypes suggest that ERK1/2 activation alone is not sufficient to induce endosomal acceleration. Engagement of alternative pathways, such as JAK-STAT by CNTF, may override ERK-mediated enhancements. Pathway bias, therefore, likely dictates effector kinase dominance and shapes motor recruitment, cargo specificity, and neuron-type selectivity (Gibbs et al., 2015; Brady & Morfini, 2017).

A further layer of regulation may stem from the balance between dynein and kinesin motor activities, as previously described (Hancock, 2014). BDNF signalling may promote a retrograde dominance by increasing dynein recruitment, reducing kinesin drag, or upregulating retrograde-specific transport proteins. Conversely, CNTF may shift the balance the other way around (e.g., by recruiting more kinesin, or reducing dynein-complexes on signalling endosomes), while non-effectors like proBDNF, HGF, and NRTN may leave it unchanged. These differences may mirror functional specialisation, with some NTFs optimised for long-distance signalling (Matusica & Coulson, 2014), and others for local plasticity, muscle homeostasis (Chevrel et al., 2006; Mills et al., 2018; Delezie et al., 2019; Correia et al., 2021) and/or sensory neuron modulation (Liebl et al., 2000).

### Receptor-Level Modulation

While this study focuses on ligands regulating endosomal transport dynamics, unravelling the complexities at the receptor level is equally important. For example, the CNTF receptor (CNTFRα) is more essential than CNTF itself, as null mutations in CNTFRα cause perinatal lethality with severe motor neuron loss, whereas CNTF knockout mice remain viable and display only mild phenotypes (DeChiara et al., 1995; Plun-Favreau et al., 2001). In mature neurons, Trk receptors act as retrograde messengers, transmitting peripheral neurotrophin signals to the soma, with evidence suggesting that activation and vesicular trafficking, rather than baseline receptor levels, govern signalling efficacy (Bhattacharyya et al., 1997). Signalling endosomes are known to carry neurotrophin receptors such as TrkB and p75^NTR^ (Deinhardt et al., 2006). Proteomic profiling of H_C_T-labelled endosomes extended these observations, detecting CNTFR and LIFR but not gp130 [CNTF], Src but not c-MET [HGF], GFRα2 but not Ret [NRTN], and p75^NTR^ with Sort1 [proBDNF] (Debaisieux et al., 2016).

Ligand-receptor complex composition is another determinant of downstream effects of NTFs. For BDNF, varying combinations of the TrkB isoforms (e.g. full length TrkB, truncated TrkB), and p75^NTR^, as homodimeric or heterodimeric complexes, drive distinct signalling outcomes (Chao, 2003; Lu et al., 2005; Meeker & Williams, 2015; Tessarollo & Yanpallewar, 2022). In this regard, we used cleavage-resistant proBDNF to selectively engage p75^NTR^-sortilin signalling pathways; however, acute stimulation with proBDNF did not alter transport dynamics. Albeit initially unexpected, this outcome is consistent with the restricted expression and role of p75^NTR^ in mature and uninjured neurons, where its functions are diminished compared to development or injury contexts (Meeker & Williams, 2015). Future studies may benefit from examining p75^NTR^-sortilin modulation in disease settings, where impaired axonal transport could unmask context-dependent effects not evident in healthy neurons.

### Additional considerations

Direct NTF influences on axonal transport can also be cargo-specific. For example, IGF1 selectively impairs signalling endosome transport while leaving mitochondrial motility unaffected, underscoring the importance of cargo identity (Fellows et al., 2020). Such effects may also extend beyond signalling endosomes via recently described “hitch-hiking” mechanisms. BDNF, for instance, enhances endosome movement yet halts mitochondrial transport through Ca^2+^-dependent interactions with Miro1, favouring local rather than long-range responses (Su et al., 2014). In addition, BDNF regulates lysosomal trafficking by promoting mTORC1 recruitment to late endosomes (Khamsing et al., 2021; Sidibe et al., 2022), potentially influencing other organelles such as RNA granules that travel associated to LAMP1-positive vesicles (Liao et al., 2019; De Pace et al., 2024). Rab7-linked alterations reinforce this principle, as they specifically impair signalling endosome trafficking without affecting lysosomal or mitochondrial transport, highlighting that cargo identity critically shapes transport outcomes (Villarroel-Campos et al., 2025b). Disentangling these multiple layers of regulation will be key to understand how NTFs differentially shape cargo trafficking and why BDNF specifically tunes the system toward efficient long-range signalling in motor neurons.

Our study focused on motor neuron axonal transport far from the injection site, but NTF effects may also manifest locally within the injected muscle, at NMJs, or in sensory neurons. Such compartmentalised actions highlight the need for integrated models accounting for spatiotemporal conditions and/or cell specificity. Crucially, signalling outcomes are influenced by receptor context, as the same ligand can act differently when engaging receptors at the soma, axon, or synapse (Bothwell, 2019). Many NTF receptors, including TrkA, TrkC, and RET, also act as dependence receptors, triggering alternative signalling when the ligand is absent (Huang & Reichardt, 2003). This complexity, together with the varied expression of NTF receptors across muscle and motor neurons, raises important questions about how receptor availability and localisation dictate the transport and signalling outcomes we observe.

### Implications for neurodegeneration/injury

NTF ligand-receptor signalling and transport recruitment are altered in neuromuscular diseases, such as amyotrophic lateral sclerosis (ALS), Charcot-Marie-Tooth disease (CMT), and following injury (Mitre et al., 2017; Stansberry & Pierchala, 2023). Acute ligand delivery, as used here, may differ markedly from chronic overexpression approaches (e.g., AAV-mediated), which can change receptor availability and feedback control. Indeed, muscle-specific BDNF gene therapy in CMT2D mice restored signalling endosome transport to wild-type levels and increased the motor adaptor Snapin in the sciatic nerve (Sleigh et al., 2023). Similarly, muscle overexpression of GDNF boosts RET expression and increased ERK/AKT signalling in the spinal cords of ALS mice, suggesting paracrine neuroprotection mediated by retrograde mechanisms (Mòdol-Caballero et al., 2021). These findings underscore that transport modulation depends on ligand identity, receptor context, and disease state, which are key considerations for therapeutic development.

## Conclusions

Our results show that NTF regulation of axonal transport is selective rather than universal. BDNF enhances transport, CNTF slows it down, and others exert no detectable effect, reflecting differences in signalling pathways, receptor interactions, and functional roles. Alterations in axonal transport are therefore highly regulated, and not a generic outcome of NTF exposure. Understanding these ligand-, receptor- and context-dependent mechanisms will be critical for designing targeted strategies to fully harness the therapeutic potential of modulating axonal transport in neurodegenerative disease and nerve repair.

## Acknowledgements

We thank the personnel of the Denny Brown Laboratories for maintaining the mouse colonies (Queen Square Institute of Neurology, University College London), and Jose Norberto Sagullo Vargas and David Villarroel-Campos (Queen Square Institute of Neurology, University College London) for critical reading of the manuscript. Schematics in the **Graphical Abstract** and **Figure 1** were generated using BioRender (https://www.biorender.com/). This work was supported by the following awards: Junior Non-Clinical Fellowship and Project Grant from the Motor Neuron Disease Association (Tosolini/Oct20/973-799) (APT); a Col Bambrick MND Research Grant from Motor Neuron Disease Research Australia (IG 2450) (APT); FightMND Drug Development Grant awarded to Giovanni Nardo (Istituto di Ricerche Farmacologiche Mario Negri - IRCCS) (DDG-73; for APT); Wellcome Trust Senior Investigator Awards (107116/Z/15/Z and 223022/Z/21/Z) (GS); UK Dementia Research Institute award (UKDRI-1005) (GS); a UK Medical Research Council (MRC) award (MR/Y010949/1) (JNS); and the UCL Therapeutic Acceleration Support scheme supported by funding from MRC IAA 2021 (UCLMR/X502984/1) (JNS).

## Author contribution

Conceptualisation: APT, JNS, GS. Investigation: ERR, APT. Data analysis: ERR, APT. Resources: APT, GS. Writing: ERR, JNS, GS, APT. Supervision: JNS, GS. Funding Acquisition: APT, JNS, GS

## Competing Interests

The authors declare no competing interests.

## Data Availability

The datasets generated during and/or analysed during the current study are available from the corresponding author on reasonable request.

## Notes

### Competing Interest Statement

The authors have declared no competing interest.

